# Amyloid-β fibrils induce calcium phosphate crystallization via catalysis of ATP hydrolysis

**DOI:** 10.1101/2025.07.12.664505

**Authors:** Sisira Mambram Kunnath, Elad Arad, Elinor Slavsky, Lonia Friedlander, Hanna Rapaport, Raz Jelinek

## Abstract

Brain-localized deposits of crystalline calcium phosphate, mainly comprising of hydroxyapatite, are a pathological hallmark of Alzheimer’s disease (AD). While calcium phosphate crystals are specifically identified in the neuronal amyloid plaques in AD, consisting mostly of beta amyloid (Aβ) fibrils, the mechanisms and factors affecting the biomineralization are unknown. Here, we present a novel mechanism for AD plaque-induced formation of crystalline calcium phosphate. Specifically, we show, for the first time, that Aβ amyloid fibrils catalyse dephosphorylation of adenosine triphosphate (ATP). Furthermore, incubating Aβ fibrils, ATP, and calcium ions gave rise to pronounced deposition of hydroxyapatite crystals upon the Aβ amyloid fibril matrix, originating through reaction between calcium ions and the monophosphate released through the catalytic dephosphorylation reaction. This pathway may explain amyloid plaque-associated calcification in AD, as elevated levels of both ATP and calcium are key features of the disease.

## Introduction

Extensive calcification in the brains of Alzheimer’s disease (AD) patients is among the hallmarks of the disease [1,2]. Calcium phosphate crystallites, mainly the hydroxyapatite (hAp) isoform, have been particularly identified as key constituents of AD calcification [3]. The mineral deposits accumulate in brain regions critical for memory and cognition, such as the hippocampus and cortex, and have been specifically observed at the amyloid plaques of AD patients [4, 5]. While the factors affecting calcium phosphate biomineralization in AD are unknown, this phenomenon has been linked to disruption of calcium homeostasis, specifically elevated calcium levels in the brain of AD patients [6,7,8]. ATP disruption has been also an underlying feature in AD [9]. Neurons in AD-impacted brains often show impaired oxidative phosphorylation, leading to energy deficits [10]. Moreover, damaged or dying neurons in AD release ATP, which activates microglia, leading to chronic neuroinflammation, a key contributor to AD pathology [11]. Some studies suggest ATP modulates Aβ aggregation and toxicity [12]. Dysregulated purinergic signalling linked to elevated extracellular ATP is believed to add another layer of neuronal stress. [13, 14]

Recent groundbreaking discoveries in our laboratory and by others have revealed that native (physiological) amyloid fibrils catalyse pathological and metabolic reactions. Specifically, we showed catalysis of key biological reactions by amyloid-beta (Aβ) fibrils [15] and glucagon amyloid fibrils [16,17]. Furthermore, we have also recently shown that phenol-soluble modulins (PSMs), prominent bacterially secreted functional amyloids, catalyse hydrolysis and degradation of β-lactam antibiotics, pointing to a possible mechanism for bacterial antibiotic resistance [18]. Alpha-synuclein fibrils were also reported to exhibit catalytic activities, including ATP hydrolysis [19, 20]. These studies highlight potentially new functional roles, yet unknown, for amyloid fibrils in Nature.

Here, we report on an intriguing Aβ-mediated pathway for calcium phosphate biomineralization. Specifically, we show, for the first time, that Aβ amyloid fibrils catalyse ATP dephosphorylation. Consequently, the monophosphate ions generated through the catalytic hydrolysis react with co-added Ca^2+^ ions, giving rise to substantial crystalline deposits of calcium phosphate, particularly hydroxyapatite, directly associated with the Aβ amyloid fibril matrix. The role Aβ amyloid fibrils as a catalyst underscores a possible route for aberrant calcium phosphate biomineralization in AD.

## Results and discussion

Figure 1. illustrates the physiological framework and hypothesis guiding this study. Specifically, we focus on the observation of ubiquitous calcium phosphate biomineralization, associated with the neuronal amyloid plaques in the brains of AD patients [3]. We hypothesize that Aβ amyloids catalyse dephosphorylation of ATP, generating monophosphate residues which react with Ca^2+^, yielding crystalline deposits of calcium phosphate. Importantly, aberrant, disregulated levels of both ATP and calcium ions are present in the brains of AD-inflicted patients [21]. Moreover, varied amyloidogenic proteins have been reported to catalyse ATP hydrolysis [18,19], although ATP dephosphorylation by Aβ amyloid fibrils have not been reported yet.

**Figure 1:**
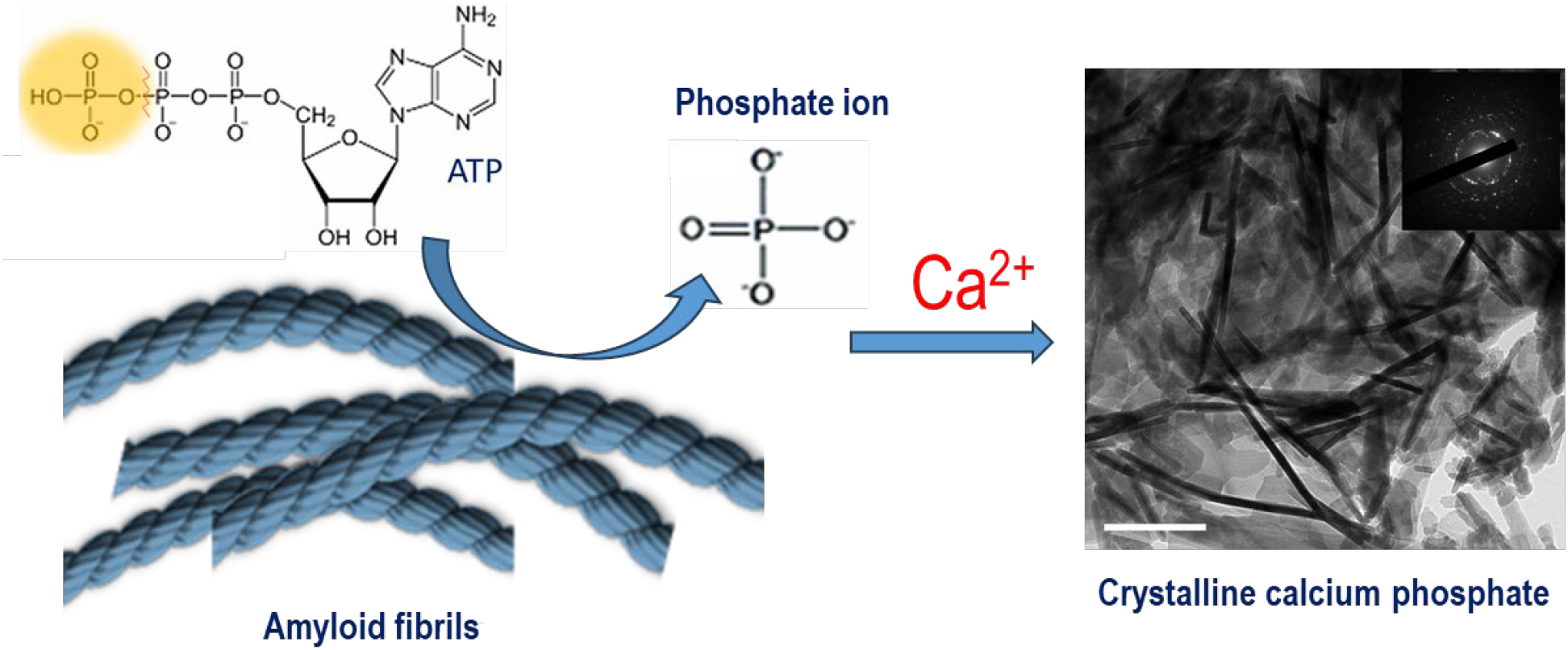
The experimental scheme depicting calcium phosphate crystallization induced through reaction between calcium and phosphate ions released by Aβ-catalysed ATP dephosphorylation.

**Figure 2**. depicts the catalytic properties of Aβ fibrils towards dephosphorylation reactions. We initially tested the impact of incubating Aβ fibrils with para-nitrophenyl orthophosphate (pNPP), a model substrate widely used to probe phosphatase activity and phospho-ester bond hydrolysis (**Figure 2a**). Specifically, pNPP dephosphorylation can be monitored via recording the visible absorbance (at 400 nm), accounting for the para-nitrophenol (pNP) hydrolysis product. Indeed, the kinetic curves in Figure 2a,i reveal that dephosphorylation of pNPP was significantly more pronounced in the presence of Aβ fibrils (blue curve), as compared to the Aβ monomer solution (red curve), or buffer conditions (no Aβ peptide). The graph depicting the initial pNPP hydrolysis reaction rate (V_0_) as a function of fibril concentration (at a constant pNPP concentration of 1 mM) features a linear relationship (Figure 2a,ii), indicative of first-order reaction kinetics [22]. Importantly, the curve of V_0_ vs pNPP substrate concentration in Figure 2a,iii reveals a hyperbolic relationship, reflecting a Michaelis–Menten mechanism, the hallmark enzymatic reaction profile [23].

**Figure 2:**
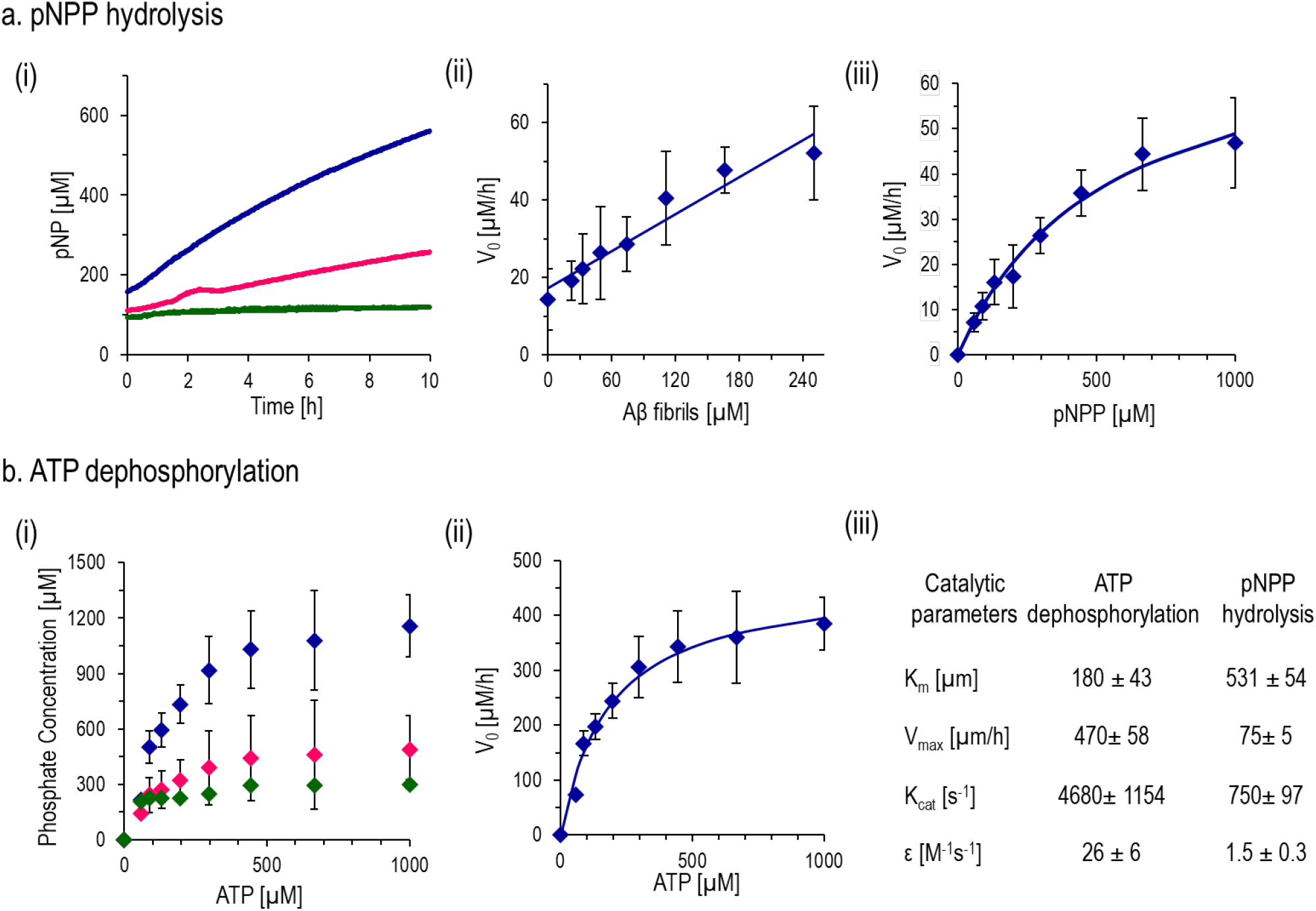
Phosphatase-like catalytic activity of Aβ fibrils. **a. pNPP hydrolysis** (i) pNPP hydrolysis recorded upon incubation with Aβ fibrils (blue curve) and Aβ monomers (pink curve) (Aβ concentration 50 μM, 50mM HEPES buffer, 7.4 pH). The concentration of hydrolysis product pNP was determined by measuring the absorbance at 400 nm in the reaction mixture. Control buffer reaction was also performed (green curve) (ii) Initial hydrolysis reaction rate (V0) as a function of Aβ concentration in the presence of pNPP (1 mM). (iii) Initial hydrolysis reaction rate as a function of pNPP concentration (Aβ concentration was 50 μM). **b. ATP dephosphorylation** (i) Phosphate product concentration recorded after 3-hr incubation (using the ATPase assay) in the presence of Aβ fibrils (blue curve), Aβ monomers (pink curve) and only buffer (20mM tris buffer, pH 8, green curve). (ii) Dephosphorylation reaction rate (V0) as a function of ATP concentration in the presence of Aβ fibrils (50 μM). (iii) Catalytic parameters corresponding to the ATP dephosphorylation and pNPP hydrolysis. The curve was fitted according to Michaelis−Menten kinetics. Results are presented as average ± SEM, n = 5.

While Figure 2a demonstrates that Aβ fibrils catalyse hydrolysis of phosphate bonds in the model substrate pNPP, we also tested whether Aβ amyloid fibrils catalyse dephosphorylation of ATP (**Figure 2b).** To assess ATP dephosphorylation, we employed the malachite green-based ATPase activity assay, yielding a quantifiable colour change (recorded at 620 nm) upon binding of the dye to PO_4_^-^ released upon ATP dephosphorylation [16]. Indeed, Figure 2b,i reveals that Aβ fibrils catalyse ATP dephosphorylation, giving rise to significantly higher phosphate concentration compared to either Aβ monomers or buffer. The rate of Aβ-induced ATP dephosphorylation (V_0_) as a function of ATP concentration yielded a hyperbolic curve (Figure 2b,ii), reflecting a Michaelis–Menten mechanism [23]. The catalytic parameters for the Aβ-mediated ATP dephosphorylation and pNPP hydrolysis, extracted through fitting the experimental parameters to the Michaelis-Menten equation, are outlined in Figure 2b,iii. Notably, the Michaelis constant (K_m_) for Aβ fibrils-catalysed ATP dephosphorylation is 180 µM, which is even lower than the value observed for pNPP, indicating higher affinity between ATP and Aβ fibrils. Figure 2b,iii also reveals for the catalytic ATP dephosphorylation a maximum reaction rate (V_max_) of 470 μM/h, resulting in a turnover number (Kcat) of 4680 s^−1^ and a catalytic efficiency (ε) of 26 M^−1^ s^−1^. Importantly, while the catalytic efficiency of the Aβ amyloid fibrils is lower than that of other ATPase-mimicking peptides, the turnover number (Kcat) is ∼ 10^4^ times lower than the natural ATPase, and the reaction rate is also slow compared to other ATPase-mimicking peptides [24], it still represents a significant catalytic activity [16]

Figure 3. presents experiments designed to illuminate the mechanistic features of Aβ fibrils-catalysed ATP dephosphorylation, specifically binding interactions between ATP and Aβ amyloid fibrils. **Figure 3a** depicts fluorescence emission spectra of a 2’,3’-o-(2,4,6-trinitrophenyl) adenosine 5’-triphosphate monolithium trisodium salt (TNP-ATP), a fluorescently labelled ATP analogue, with and without co-added Aβ fibrils. Notably, the fluorescence emission spectrum shows a strong peak for TNP-ATP at 550 nm. Notably, in the presence of Aβ fibrils, the peak intensity of TNP-ATP at 550 nm significantly increased (together with a minute red shift) is likely ascribed to proximity between Aβ and TNP-ATP, giving rise to quenching of the fluorophore [25]. ATP-induced Aβ autofluorescence quenching (Figure S1) is consistent with this interpretation.

**Figure 3:**
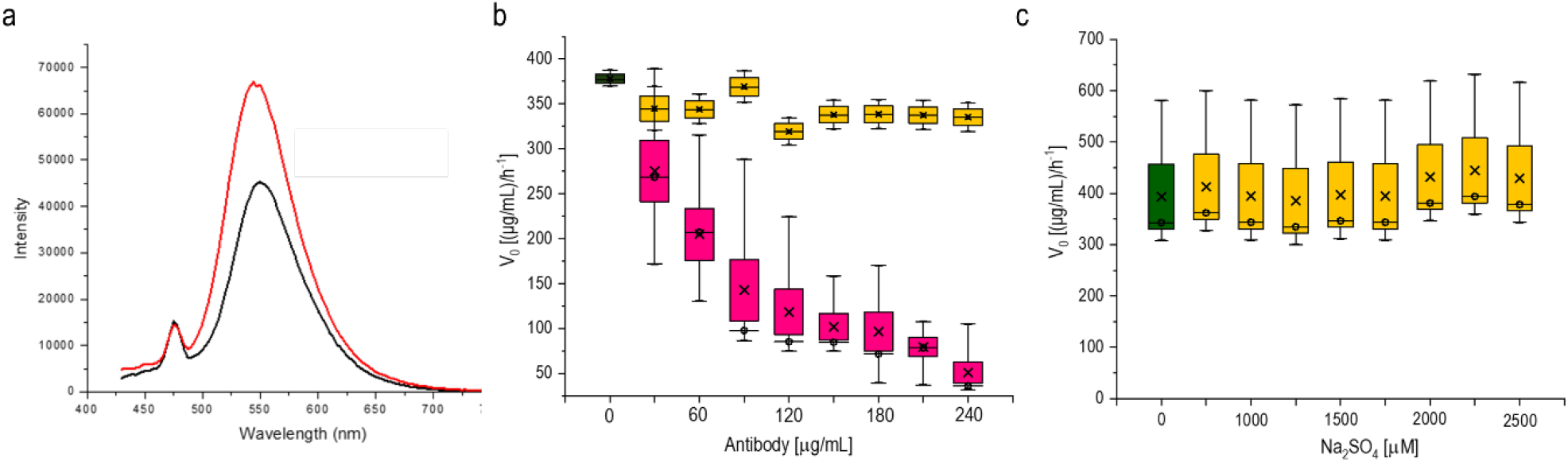
Analysis of ATP / Aβ fibrils interactions. ALL LETTERS SHOULD BE BIGGER. **a.** Fluorescence emission spectra of TNP-ATP alone (2mM, black) and TNP-ATP in the presence of Aβ fibrils (50μM, red), excitation at 410 nm **b.** Effect of an anti-ATPase antibody on Aβ fibril-catalysed ATP dephosphorylation. Box-and-whisker plot depicting the initial ATP dephosphorylation reaction rates in the presence of Aβ fibrils (50μM), upon addition of different concentrations of a specific antibody (Anti ATPase antibody; pink boxes), and a non-specific polyclonal antibody [HXK I antibody (N-19); yellow boxes]. The green box represents the ATP-Aβ fibrils sample without an antibody added. Data depict the mean line and quartile calculations using inclusive median. The measurements were repeated three times in independent experiments (n=3) for anti ATPase and five times in independent experiments (n=5) for HXK I antibody (N-19). **c.** Effect of SO_4_ ^2-^ upon Aβ fibril-catalysed ATP dephosphorylation. A box-and-whisker plot depicting the ATP dephosphorylation reaction rates in the presence of Aβ fibrils (50μM), upon addition of different concentrations of Na2SO4 (yellow boxes). The green box represents the ATP-Aβ fibrils sample in buffer. Data show mean line and quartile calculation using inclusive median. The measurements were repeated three times in independent experiments (n=3)

To assess whether presumptive ATP-specific catalytic sites are present on the fibrils’ surface, we tested the effect of an anti-ATPase antibody on the reaction rate (**Figure 3b**). Notably, the box-and-whisker plot in **Figure 3b** shows that co-addition of the anti-ATPase antibody to the reaction mixture (at a significantly lower concentration compared to Aβ) effectively inhibited the catalytic activity of the Aβ amyloid fibrils in a concentration-dependent manner. As a control, we also assessed the impact of co-incubating Aβ amyloid fibrils and ATP with a non-specific polyclonal antibody, human hexokinase I (HXK I, Figure 3b, yellow boxes). Indeed, Figure 3b demonstrates that HXK-I did not inhibit Aβ fibrils-induced catalytic activity. Overall, the catalysis data in Figure 3b underscore the specificity of the Aβ amyloid fibrils towards ATP and the apparent presence of ATPase-mimetic sites upon the Aβ fibrils’ surface.

Yet in another control experiment, presented in **Figure 3c**, we examined whether the catalytic activity of Aβ fibrils towards ATP dephosphorylation is driven by electrostatic attraction between the fibrils and the negatively charged ATP. In the experiment, we tested the effect of co-adding to the Aβ fibrils and ATP mixture SO_4_^2-^, exhibiting double negative charge and previously employed as a mimic for ATP [26], Notably, the presence of varying concentrations of SO42-anions in the solution did not display inhibitory effect upon the catalytic activity of Aβ amyloid fibrils, underscoring the specificity of the fibrils towards ATP.

While Figures 2 and 3 attest to efficient catalysis of ATP dephosphorylation by Aβ amyloid fibrils and binding specificity of ATP towards the fibrils, we further investigated whether the release of phosphate residues following the hydrolysis reaction leads to calcium phosphate biomineralization, upon co-addition of Ca^2+^ ions (**Figure 4**). In the experiment, we co-added 5mM CaCl2 to the reaction mixture of ATP and Aβ fibrils (20mM) at (the physiological) 7.4 pH. The electron microscopy results in **Figure 4a-b** furnish remarkable visual evidence for the occurrence of Aβ amyloid fibril-induced calcium phosphate biomineralization. The scanning electron microscopy (SEM) images in Figure 4a (i-iii), capturing the reaction mixture after different incubation times, demonstrate accumulation of substantial deposits upon the fibrils’ matrix. The SEM images in Figure 4b underscores that significant deposition of seemingly crystalline aggregates occurred only in the presence of Aβ fibrils, ATP, and calcium ions; no deposits were observed without Aβ fibrils present. The representative electron dispersion spectrum (EDS) in **Figure 4c** confirms that the deposits comprise of calcium phosphate; notably, the Ca/P ratio obtained - 1.68 - indicates the formation of hydroxyapatite, the dominant calcium phosphate crystalline isoform in AD [27].

**Figure 4:**
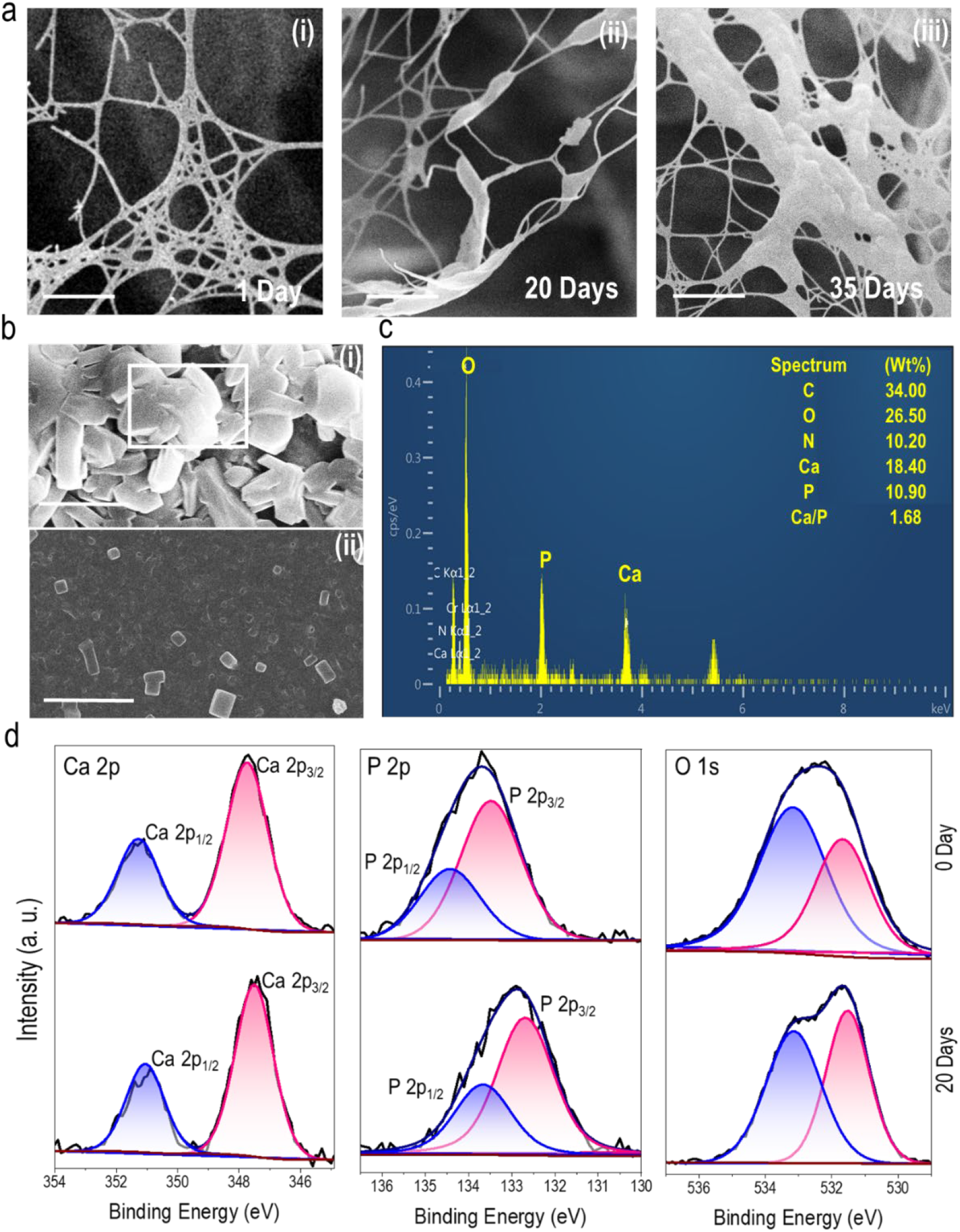
Calcium phosphate biomineralization upon incubation of Aβ fibrils, ATP, and calcium ions. **a.**SEM images recorded at different incubation times. Pre-formed Aβ fibrils (50μM, in 20mM tris buffer, pH 8) co-incubated with ATP (2mM) for 3h at 37°C and subsequent addition of CaCl2 (5 mM). The scale bar is 5μm. **b.** SEM images recorded after 35-day incubation with (top) and without (bottom) Aβ fibrils in the reaction mixture. Scale bar is 100 μm. **c.** EDS acquired in the region indicated by the square in panel b. **d.** XPS of the reaction mixture recorded immediately after co-addition of Aβ fibrils, ATP, and calcium ions (top spectra) and after 20-day incubation.

The XPS data in **Figure 4d** corroborate the SEM and EDS results, attesting to formation of hydroxyapatite deposits in the reaction mixture comprising Aβ fibrils, ATP, and calcium chloride. Specifically, the Ca 2p3/2 binding energy shifts from 347.72 eV initially to 347.52 eV after 20-day incubation, consistent with hydroxyapatite biomineralization [28]. Similarly, the P 2P3/2 peaks shifted from 133.47 eV to 132.68 eV after 20-day incubation, reflecting the transformation of the phosphate from the organic phosphate species in ATP into hydroxyapatite [29,30]. The oxygen XPS result similarly accounts for a significant structural transformation associated with calcium phosphate biomineralization. The O 1s spectrum recorded initially (0 days) accounts for the organic phosphate in ATP as well as the –OH groups. After 20-day incubation, the intensity of the peak at 531.8 eV increases, as well as a shifting to 531.5 eV was recorded, consistent with the P–O bond in hydroxyapatite (HA) [28].

Figure 5. depicts diffraction data confirming the formation of crystalline hydroxyapatite deposits upon incubation of Aβ fibrils, ATP, and calcium chloride. The X ray diffraction (XRD) patterns in **Figure 5a** reveal the appearance of the predominant hydroxyapatite crystal planes at (100), (200), (211), and (203) [corresponding to 2θ values 10.8, 21.8, 31.8, and 45.0, respectively [31] after 20-day incubation. These peaks are absent following the initial mixing of the reaction constituents (only the (200) NaCl plane is apparent in the XRD pattern recorded for this sample, Figure 5a).

**Figure 5:**
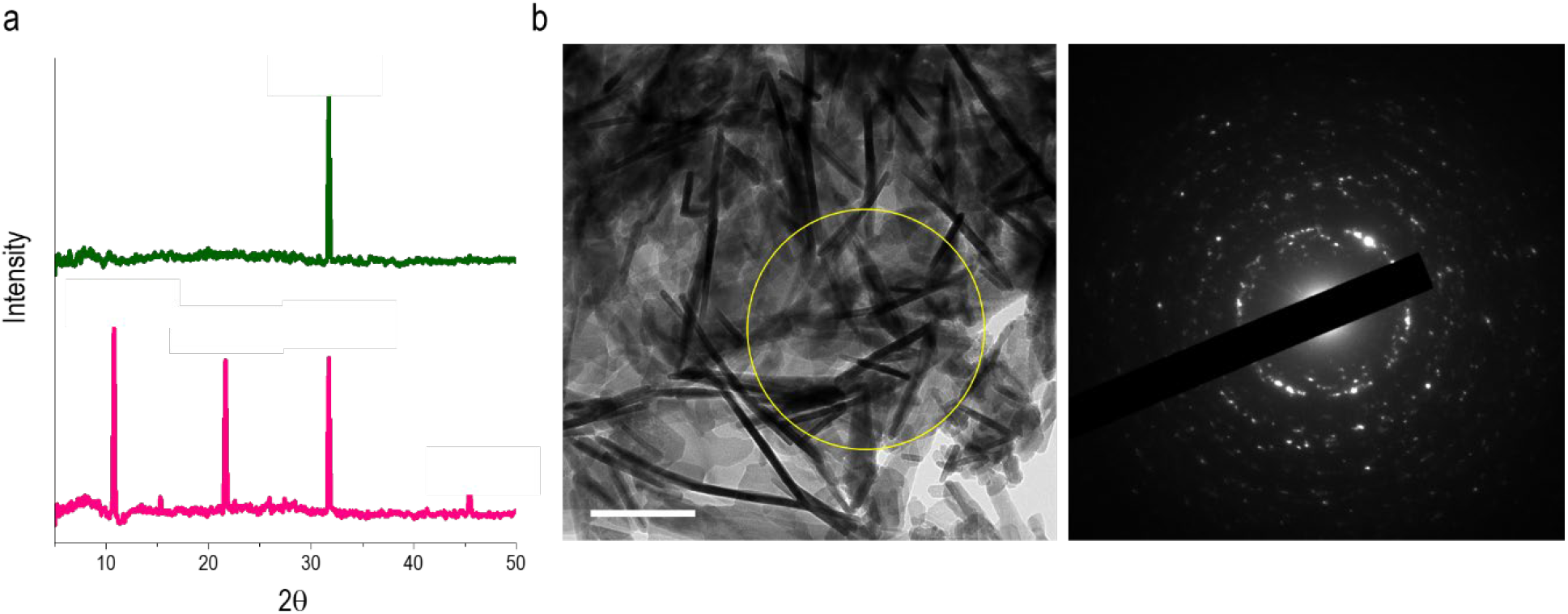
Formation of crystalline hydroxyapatite after incubation of Aβ fibrils, ATP, and Ca2+ ions. **a.** XRD spectra of the reaction mixture represents the pink plot (20 days of reaction) and the 0 days of the reaction represents the green plot in the graph. **b.** HRTEM image showing the crystalline deposits after 35-day incubation. Scale bar is 200nm. SAED pattern from the marked yellow region of the HRTEM sample on the right.

The representative HRTEM image in **Figure 5b,** recorded after 20-day incubation, clearly reveal the assembly of calcium phosphate crystallites. The bright diffraction spots apparent in the selected area electron diffraction (SAED) pattern further underscores the formation of crystalline hydroxyapatite (the specific crystal planes corresponding to the diffraction spots are indicated in Figure S2). The SAED pattern complements the XRD data in Figure 5a, as the multitude of crystal planes observed reflect crystalline nature of the sample, evidenced by distinct diffraction spots corresponding to various lattice planes ((100), (200), (211) and (203). These features confirm the presence of well-ordered crystalline domains and are consistent with the phase assignments derived from the XRD pattern.

## Conclusions

We report on a novel molecular pathway by which Aβ fibrils may induce calcium phosphate biomineralization, a pathological hallmark of Alzheimer’s disease. Specifically, we demonstrate that Aβ fibrils catalyze dephosphorylation of ATP, resulting in the release of inorganic phosphate. In the presence of calcium ions, nucleation and growth of hydroxyapatite crystals occurred on the amyloid fibril matrix. The deposition of calcium phosphate crystallites depended solely on the presence of Aβ fibrils, which display specific phosphatase-like catalytic sites responsible for ATP docking and induction of the dephosphorylation reaction. This remarkable mechanism may provide a biochemical basis for amyloid plaque-associated calcification in AD, especially considering the aberrant elevated levels of both ATP and calcium ions associated with the disease. This novel Aβ-mediated calcium phosphate biomineralization pathway may usher new understanding of AD pathologies and aid the search for potential therapeutic avenues.

### Experimental section

#### Materials

Human Amyloid β (residue 1-42), of molecular weight 4514.04 g/mol, exhibiting the sequence DAEFRHDSGYEVHHQKLVFFAEDVGSNKGAIIGLMVGGVVIA, >95% purity was obtained as lyophilized powders from AnaSpec, Inc. (Fremont, USA), HEPES (4-(2-hydroxyethyl)-1-piperazineethanesulfonic acid, >99% purity) was obtained from Holland-Moran (Yehud, Israel). 1,1,1,3,3,3-Hexafluoro-2-propanol (HFIP, 99% purity) and 4-nitrophenyl-disodium orthophosphate (pNPP) were purchased from Sigma-Aldrich (Rehovot, Israel). Adenosine 5′-triphosphate disodium salt trihydrate (ATP) (Aaron Chemicals, USA), Tris(hydroxymethyl)aminomethane, ACS, 99.8-100.1% (Thermo Scientific Chemicals, USA), Calcium Chloride dihydrate 99% pure (Thermo Scientific–ACROS, USA), ATPase/GTPase Activity Assay Kit -catalog number-MAK113-1KT (Sigma-Aldrich, Israel), Anti-Na+/K+ ATPase α-1 Antibody, clone C464.6 (Sigma-Aldrich, Germany), A nonspecific antibody HXK I Antibody (N-19), catalog number sc-6517, isotype: IgG1(Santa Cruz Biotechnology, Santa Cruz, California). Sodium sulfate, reagent plus TM, 99.0% (Sigma-Aldrich (Rehovot, Israel)

#### Peptide sample preparation

Aβ (1-42) peptides were initially dissolved in 1,1,1,3,3,3-hexafluoro-2-propanol (HFIP) and titrated with 30% ammonium hydroxide until the solution was optically clear, ensuring complete solubilization and minimizing pre-aggregation. Stock solutions (10 mM) were prepared and stored at −20 °C. Prior to each experiment, aliquots were thawed, transferred into glass tubes, and subjected to vacuum evaporation (0–20 mbar, 3 h) to remove residual HFIP. The resulting dry peptide films were reconstituted in the appropriate buffer. All buffers were prepared using deionized water (18.2 MΩ·cm; Barnstead Smart2Pure, Thermo Scientific). Peptide solutions were incubated at 37 °C for 24 h with gentle agitation (150–200 rpm) to ensure dispersion and equilibration prior to use.

#### pNPP hydrolysis

The hydrolysis of pNPP was assessed by tracking the formation of para-nitrophenol (pNP) over a 10-hour period. A 1000 μM solution of pNPP was freshly prepared in 50 mM HEPES buffer (pH 7.4) immediately before each experiment and subsequently diluted 2/3 fold in the same buffer. To initiate the reaction, 140 μL of the diluted pNPP solution was mixed with 70 μL of preincubated Aβ (50 μM in 50 mM HEPES, pH 7.4). The reaction mixtures (60 μL per well) were promptly dispensed in triplicate into a 384-well flat-bottom clear microplate (Greiner, F-bottom), sealed with optical adhesive film, and gently agitated for 30 s. Kinetic measurements were performed using a Multiskan GO microplate reader (Thermo Fisher Scientific, Waltham, MA) pre-equilibrated to 37 °C. Absorbance at 400 nm was recorded at one minute intervals for a total duration of 10h to quantify pNP production.

#### ATP dephosphorylation

ATP dephosphorylation was monitored by quantifying the formation of inorganic monophosphate over a 3-hour incubation period. Fresh aqueous ATP solution (20 mM) was prepared immediately prior to each experiment. Reactions were initiated by combining preformed Aβ fibrils with ATP in 20 mM Tris buffer (pH 8.0), followed by incubation at 37 °C for 3 h. At the end of the reaction, 165 μL of the reaction mixture was quenched by the addition of 55 μL malachite green reagent in the ATP/GTPase assay kit, and the samples were further incubated for 30 min at room temperature to allow colour development. Phosphate release was quantified by measuring absorbance at 620 nm using a Multiskan GO microplate reader (Thermo Fisher Scientific, Waltham, MA). Measurements were performed in triplicate using 60 μL per well in a 384-well flat-bottom clear plate (Greiner, F-bottom). Phosphate concentrations were interpolated from a standard curve (0–50 μM), and reaction rates were calculated by dividing the amount of free phosphate by the total incubation time (3 h).

#### Kinetic Data Analysis

Initial reaction velocities (V_0_) for pNP production were determined from the linear portion of the absorbance curves, corresponding to the first 45 minutes of the reaction. These initial rates were then fitted using nonlinear regression in Microsoft Excel (Solver add-in), applying the Michaelis–Menten equation:

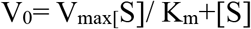

#### Kinetic Analysis of Anti-ATPase Antibody Inhibition

For kinetic assessment of the antibody effect, 20 µL of pre-incubated Aβ fibrils (final concentration: 75 μM) were dispensed into wells of a clear 384-well flat-bottom plate (Greiner Bio-One). 20 µL of anti-ATPase antibody (stock concentration: 100 µg/mL) were then added and serially diluted in a 2/3 ratio across the plate to assess dose-dependent effects. Prior to measurement, the plate was placed on 37°C for 30 minutes. Subsequently, 50 µL of freshly prepared ATP (1 mM final concentration) were added to each well and incubated at 37°C for 3h. At the end of the reaction, 20 μL of the reaction mixture was quenched by the addition of 5 μL malachite green reagent in the ATP/GTPase assay kit, and the samples were further incubated for 30 min at room temperature to allow colour development. Phosphate release was quantified by measuring absorbance at 620 nm using a Multiskan GO microplate reader (Thermo Fisher Scientific, Waltham, MA). Phosphate concentrations were interpolated from a standard curve (0–50 μM), and reaction rates were calculated by dividing the amount of free phosphate by the total incubation time (3 h). For a negative control we performed the same experiment with a non-specific polyclonal antibody, human hexokinase I (HXK I). Additional control was carrying out the experiment without antibodies co-added.

#### Competition experiments

To examine whether phosphate-like anions can compete with ATP, Na_2_SO_4_ was added in increasing concentrations to the Aβ fibrils and ATP reaction mixture. For kinetic assessment of the Na2SO4 effect, 20 µL of pre-incubated Aβ fibrils (final concentration: 75μM) were dispensed into wells of a clear 384-well flat-bottom plate (Greiner Bio-One). Subsequently, 100 µL of Na2SO4 (stock concentration: 5mM) were added and serially diluted in a 2/3 ratio across the plate to assess dose-dependent effects.

#### Scanning Electron Microscopy (SEM)

10μL samples were drop-casted on a 1cm^2^ cover glass and allowed to dry at room temperature overnight. The samples were coated with 20nm chromium to prevent the charging during the imaging. The Thermo Fisher Verios 460L field-emission scanning electron microscope (FESEM) equipped with an energy dispersive spectroscopy (EDS) used for imaging the sample.

#### X-ray photoelectron spectroscopy (XPS)

10 μL samples were drop-casted on a silicon wafer and allowed to dry overnight at room temperature. X-ray photoelectron spectroscopy (XPS) characterization was performed using an “ESCALAB Xi+” by Thermo-Fisher scientific (UK). The ambient pressure in the chamber is <2·10-10 mbar. Photoelectron emission is achieved using an Al Kα radiation source (1486.68eV) at room temperature. The spectra is collected at 90° from the X-ray source. A low-energy electron flood gun is used to minimize charging at the surface. Measurements are taken with a spot diameter of 650µm. Avantage software version 6.6.0. by Thermo-scientific was used for the XPS spectra processing. A high-resolution spectrum was measured for each relevant element along with a survey spectrum. Calibration was performed using the C1s spectra peak at 284.8eV.

#### X-Ray Diffraction (XRD)

Samples were drop-cast onto watch slide cover slips and dried to extract any present crystalline phases from solution. Dried samples were measured by thin-film X-ray diffraction (XRD) in reflectance geometry, using a symmetric (gonio) scan measurement in the 2θ range 5 - 60° over the range of the characteristic peaks of biologically relevant Ca-phosphate minerals. The data were collected on a Panalytical Empyrean III multi-purpose diffractometer (Cu Ka radiation, 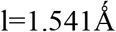) equipped with a pixCEL 3D detector in 1D scanning line-detector mode, and with an X-ray tube operated at v=45 kV, I = 40 mA. The instrument was outfitted with Panalytical multi-core™ automated optics and a 3-axis eulerian (χ-Φ-z) cradle for fine sample alignment. The sample was mounted onto a zero-background pure Si plate, which was placed directly on the eulerian cradle. Data interpretation and phase analysis was conducted using the HighScore Plus (version 5.1) XRD analysis software suite, in combination with the International Center for Diffraction Data (ICDD) PDF-5+ (2025) diffraction library

#### High Resolution Transmission Electron Microscopy (HRTEM)

After adding CaCl2 at different time points 10μL of the reaction and control samples were mixed in 90μL of methanol and mixed well. 3μL solution was drop-casted on Ultrathin Lacey carbon on gold grid of mesh 300. The samples were allowed to dry in the room temperature and imaged using Analytical High Resolution TEM-JEOL JEM 2100F with an electron source: Schottky field-emission gun operating at accelerating voltages operated at an accelerating voltage of 200 kV and energy spread of 0.8 eV.

## Supporting Information

**Figure S1:**
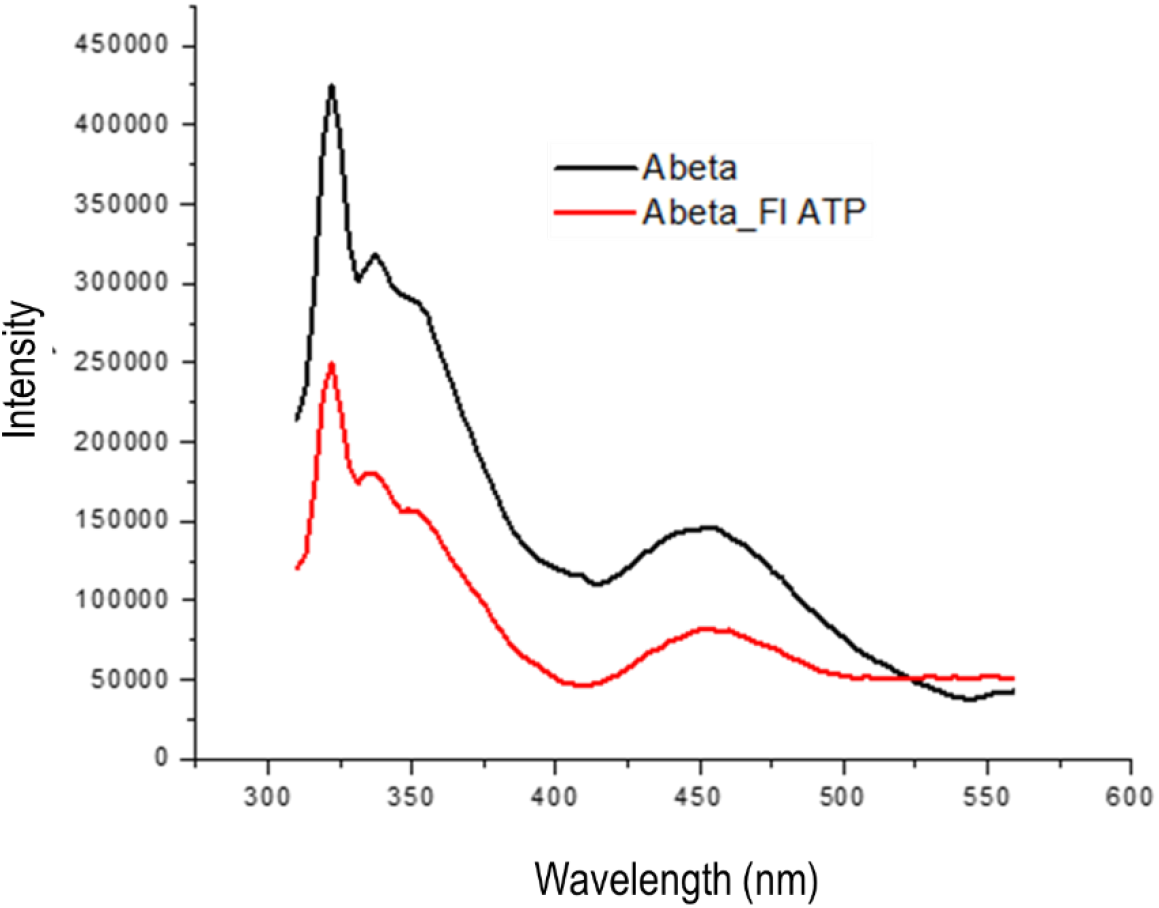
Autofluorescence of Aβ fibrils with and without ATP.

**Figure S2:**
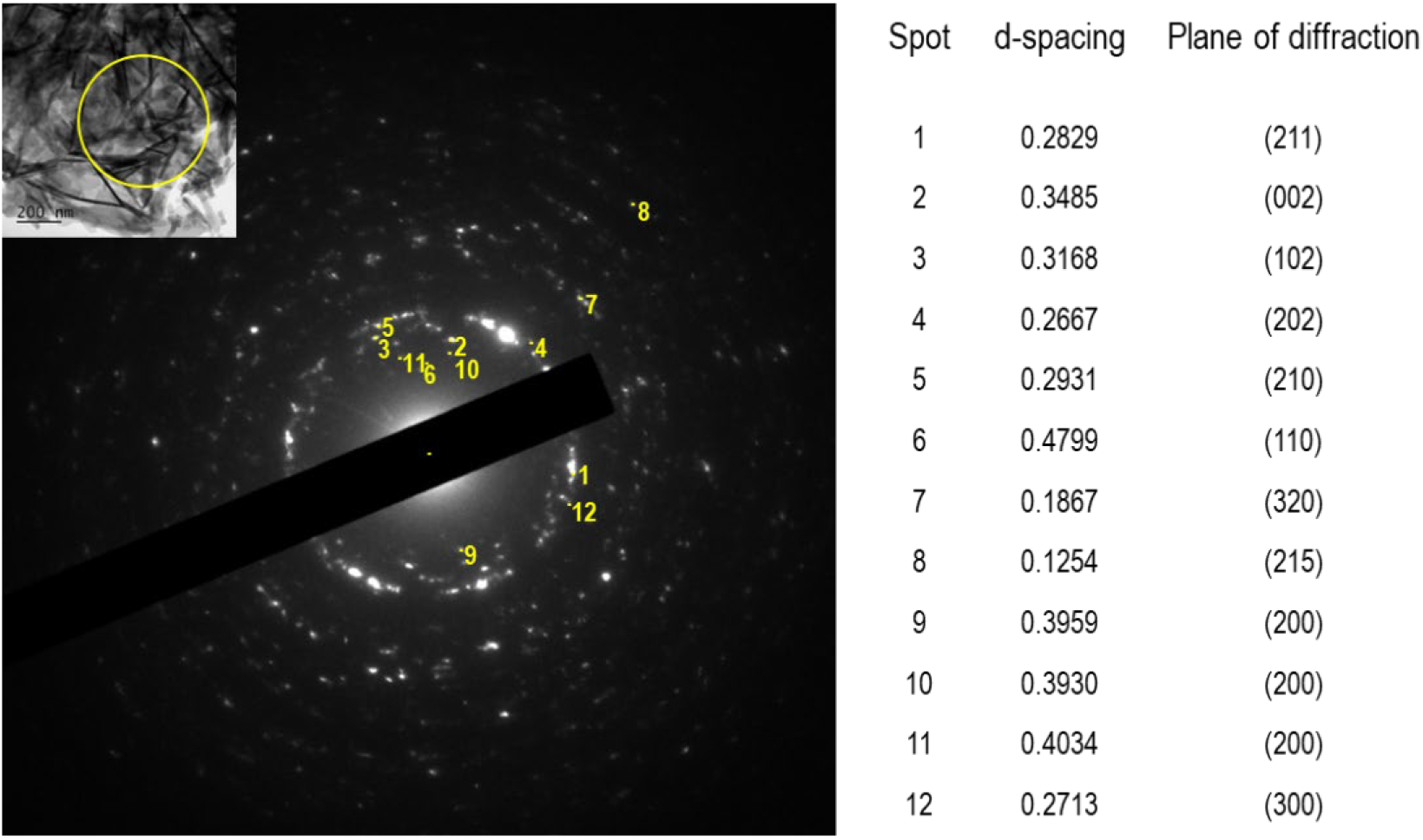
calcium phosphate crystal planes ascribed to the diffraction spots in the SAED pattern.

## References

1. Fujita, D., Terada, S., Ishizu, H., Yokota, O., Nakashima, H., Ishihara, T., & Kuroda, S. (2003). Immunohistochemical examination on intracranial calcification in neurodegenerative diseases. Acta neuropathologica, 105, 259–264.

2. Lemos, R. R., Ferreira, J. B. M. M., Keasey, M. P., & Oliveira, J. R. (2013). An update on primary familial brain calcification. International review of neurobiology, 110, 349–371.

3. Tsolaki, E., Csincsik, L., Xue, J., Lengyel, I., & Bertazzo, S. (2022). Nuclear and cellular, micro and nano calcification in Alzheimer’s disease patients and correlation to phosphorylated Tau. Acta Biomaterialia, 143, 138–144.

4. Wegiel, J., Kuchna, I., Wisniewski, T., De Leon, M. J., Reisberg, B., Pirttila, T., … & Lehtimaki, T. (2002). Vascular fibrosis and calcification in the hippocampus in aging, Alzheimer disease, and Down syndrome. Acta neuropathologica, 103, 333–343.

5. Borges, K. A., Lombardi, I., Sivilli, M., Aabye, J., Romano, I., Nasruddin, S. A., … & Savinova, O. V. (2025). Brain microvascular calcification is increased in human donors with dementia compared to elderly controls: a pilot study. Frontiers in Aging Neuroscience, 17, 1557625.

6. Mattson, M. P., & Chan, S. L. (2001). Dysregulation of cellular calcium homeostasis in Alzheimer’s disease: bad genes and bad habits. Journal of Molecular Neuroscience, 17, 205–224.

7. Demuro, A., Parker, I., & Stutzmann, G. E. (2010). Calcium signaling and amyloid toxicity in Alzheimer disease. Journal of Biological Chemistry, 285(17), 12463–12468.

8. Supnet, C., & Bezprozvanny, I. (2010). The dysregulation of intracellular calcium in Alzheimer disease. Cell calcium, 47(2), 183–189.

9. Patro, S., Ratna, S., Yamamoto, H. A., Ebenezer, A. T., Ferguson, D. S., Kaur, A., … & Solesio, M. E. (2021). ATP synthase and mitochondrial bioenergetics dysfunction in Alzheimer’s disease. International journal of molecular sciences, 22(20), 11185.

10. Alqahtani, T., Deore, S. L., Kide, A. A., Shende, B. A., Sharma, R., Chakole, R. D., … & Ghosh, A. (2023). Mitochondrial dysfunction and oxidative stress in Alzheimer’s disease, and Parkinson’s disease, Huntington’s disease and amyotrophic lateral sclerosis-an updated review. Mitochondrion, 71, 83–92.

11. Lue, L. F., Kuo, Y. M., Beach, T., & Walker, D. G. (2010). Microglia activation and anti-inflammatory regulation in Alzheimer’s disease. Molecular neurobiology, 41, 115–128.

12. Pal, S., & Paul, S. (2019). ATP Controls the aggregation of Aβ16–22 peptides. The Journal of Physical Chemistry B, 124(1), 210–223.

13. Du, H., Guo, L., & Yan, S. S. (2012). Synaptic mitochondrial pathology in Alzheimer’s disease. Antioxidants & redox signaling, 16(12), 1467–1475.

14. da Silva, G. S. S., Melo, H. M., Lourenco, M. V., e Silva, N. M. L., De Carvalho, M. B., Alves-Leon, S. V., … & De Felice, F. G. (2017). Amyloid-β oligomers transiently inhibit AMP-activated kinase and cause metabolic defects in hippocampal neurons. Journal of Biological Chemistry, 292(18), 7395–7406.

15. Arad, E., Leshem, A. B., Rapaport, H., & Jelinek, R. (2021). β-Amyloid fibrils catalyze neurotransmitter degradation. Chem Catalysis, 1(4), 908–922.

16. Arad, E., Yosefi, G., Kolusheva, S., Bitton, R., Rapaport, H., & Jelinek, R. (2022). Native glucagon amyloids catalyze key metabolic reactions. ACS nano, 16(8), 12889–12899.

17. Nandi, A., Zhang, A., Arad, E., Jelinek, R., & Warshel, A. (2024). Assessing the catalytic role of native glucagon amyloid fibrils. ACS catalysis, 14(7), 4656–4664.

18. Arad, E., Pedersen, K. B., Malka, O., Mambram Kunnath, S., Golan, N., Aibinder, P., … & Jelinek, R. (2023). Staphylococcus aureus functional amyloids catalyze degradation of β-lactam antibiotics. Nature Communications, 14(1), 8198.

19. Castillo-Caceres, C., Nova, E., & Diaz-Espinoza, R. (2025). Alpha-synuclein amyloids catalyze the degradation of ATP and other nucleotides. bioRxiv, 2025–04.

20. Parate, S., Buratti, F., Eriksson, L. A., & Wittung-Stafshede, P. (2025). In silico Identification of Substrate Binding Sites in Type-1A α-Synuclein Amyloids. Biophysical Journal.

21. Popugaeva, E., Pchitskaya, E., & Bezprozvanny, I. (2018). Dysregulation of intracellular calcium signaling in Alzheimer’s disease. Antioxidants & redox signaling, 29(12), 1176–1188.

22. Rufo, C. M., Moroz, Y. S., Moroz, O. V., Stöhr, J., Smith, T. A., Hu, X., … & Korendovych, I. V. (2014). Short peptides self-assemble to produce catalytic amyloids. Nature chemistry, 6(4), 303–309.

23. Johnson, K. A., & Goody, R. S. (2011). The original Michaelis constant: translation of the 1913 Michaelis–Menten paper. Biochemistry, 50(39), 8264–8269.

24. Castillo-Caceres, C., Duran-Meza, E., Nova, E., Araya-Secchi, R., Monasterio, O., & Diaz-Espinoza, R. (2021). Functional characterization of the ATPase-like activity displayed by a catalytic amyloid. Biochimica et Biophysica Acta (BBA)-General Subjects, 1865(1), 129729.

25. Woodbury, D. J., Whitt, E. C., & Coffman, R. E. (2021). A review of TNP-ATP in protein binding studies: benefits and pitfalls. Biophysical Reports, 1(1).

26. Dumas, C., Lascu, I., Morera, S., Glaser, P., Fourme, R., Wallet, V., … & Janin, J. (1992). X-ray structure of nucleoside diphosphate kinase. The EMBO Journal, 11(9), 3203–3208.

27. Stern, A. M., Liu, L., Jin, S., Liu, W., Meunier, A. L., Ericsson, M., … & Selkoe, D. J. (2022). A calcium-sensitive antibody isolates soluble amyloid-β aggregates and fibrils from Alzheimer’s disease brain. Brain, 145(7), 2528–2540.

28. Negrila, C. C., Predoi, M. V., Iconaru, S. L., & Predoi, D. (2018). Development of zinc-doped hydroxyapatite by sol-gel method for medical applications. Molecules, 23(11), 2986.

29. He, G., Dahl, T., Veis, A., & George, A. (2003). Nucleation of apatite crystals in vitro by self-assembled dentin matrix protein 1. Nature materials, 2(8), 552–558.

30. McLeod, K., Kumar, S., Dutta, N. K., Smart, R. S. C., Voelcker, N. H., & Anderson, G. (2010). X-ray photoelectron spectroscopy study of the growth kinetics of biomimetically grown hydroxyapatite thin-film coatings. Applied surface science, 256(23), 7178–7185.

31. JCPDS file NO. 09-0432. Hydroxyapatite phase

